# Integrating Mathematical and Mouse Models Identifies T Regulatory Cell Influx as A Key Determinant of Acquired Resistance to PD-1 Immunotherapy

**DOI:** 10.1101/2025.10.31.685923

**Authors:** Rachel S Sousa, Shannon N Geels, Claire Murat, Alexander Moshensky, S Armando Villalta, John S Lowengrub, Francesco Marangoni

## Abstract

The immune system can eradicate cancer, but various immunosuppressive mechanisms active within a tumor curb this beneficial response. However, unraveling the effects of multimodal interactions between tumor and immune cells and their contributions to tumor control using an experimental approach alone is time- and resource-intensive. To identify the critical immunological features associated with tumor control and escape, we built a mechanistic mathematical model of the interactions between CD8^+^ T cells, Tregs, DCs, and tumor cells deeply rooted in current biological concepts. A distinguishing feature of our model is that it captures Treg accrual occurring after checkpoint blockade immunotherapy. After successfully fitting the model to experimental data of a mouse model of immunogenic melanoma, we generated hundreds of parameter sets, each representing a unique ‘virtual mouse’, that fit the data equally as well to capture variability across individuals. Our model indicates that the tumor and immune states before therapy are a key limiting factor of the immune response. Increasing the initial number of tumor-killing CD8^+^ T cells alone doesn’t always result in a better outcome; instead, the model implies that there exist optimal initial ratios of immune cells that will result in improved tumor control. The model further predicts that the Treg influx into the tumor is a key determinant of resistance to PD-1 immunotherapy. We validated this predictions experimentally. Overall, this integrated approach of modeling and experimental validation identified crucial determinants of resistance to immunotherapy and can be used to guide the development of more effective therapeutic strategies.

## INTRODUCTION

Checkpoint blockade immunotherapy (CBI) revolutionized the management of advanced cancer patients. CBI aims to remove the negative feedback mechanisms (checkpoints) that evolved to prevent autoimmunity yet also curb antitumor immunity. In metastatic melanoma, PD-1 blockade resulted in 32% progression-free patients three years after diagnosis, and its combination with αCTLA-4 raises this percentage to 39%.^1^ Crucially, benefits are durable: the patients that were progression-free at the three-year time point had >95% probability of being alive after 10 years.^1^ CBI is now approved to treat more than 20 clinical indications.

Despite these successes, 69-77% of patients progress within 10 years from the start of CBI.^1^ While therapy failure can be due to mutations in tumor cells that make them refractory to treatment, another leading cause of therapy resistance is the occurrence of immunosuppressive reactions within the tumor environment.^2^ “CBI-mediated immunosuppression” is an emerging concept emphasizing that, in addition to triggering anti-tumor immunity, CBI also unleashes immunosuppressive reactions that may decrease efficacy. Alternative checkpoint receptors including Tim-3 and Lag3 have been studied as immunosuppressive adaptations to CBI,^3^ paving the way for their blockade to improve the therapeutic efficacy of PD-1 blockade.^4,5^ Macrophages are the most abundant cell type in the tumor environment, and increased interferon-gamma (IFNγ) availability after CBI instructs pro-inflammatory functions that oppose tumor growth. Nonetheless, the blockade of the PD-1 / PD-L1 axis also triggers immunosuppressive responses in tumor-associated macrophages, including the expression of IL-6 and IDO.^6,7^ However, the most studied immune adaptation to CBI is the enhancement of CD4^+^FOXP3^+^ T regulatory cells (Tregs). Seminal papers from Drs. Sakaguchi and Nishikawa suggested that PD-1 blockade directly increases the proliferation and immunoregulatory function of tumor Tregs by enhancing TCR signaling.^8,9^ Moreover, our group recently showed that, in immunogenic melanoma, Treg accumulation after PD-1 blockade was due to IL-2 secretion by tumor-associated CD8^+^ T cells.^10^ Notably, CTLA-4 blockade also results in the accrual of tumor-associated Tregs in melanoma patients.^11^ Mechanistically, αCTLA-4-mediated tumor Treg accrual is due to increased co-stimulation during cognate interactions between Tregs and dendritic cells (DCs).^12^ The extensive investigations of the antitumor immune response continue to provide new mechanistic details of the complexities of the tumor-immune system. Thus, we reason that unraveling the effects of multimodal interactions between tumor and immune cells and their contributions to tumor control using an experimental approach alone would rapidly exceed the available time and resources. To expedite and guide experimental discovery, we built a mechanistic mathematical model of the interactions between CD8^+^ T cells, Tregs, DCs, and tumor cells. We specifically built the model so that we could study antitumor responses as well as CBI-mediated immunosuppression. Our simulations indicated that tumor growth was profoundly impacted by the number of tumor cells that engrafted at early time-points and by complex alterations in the individual immune populations. We verified these predictions in vivo by transferring graded numbers of tumor cells and by analyzing mice bearing the natural microbiota that primes the antitumor immune response. Crucially, when we simulated PD-1 blockade, we identified Treg influx as a key parameter contributing to acquired resistance, and we validated this prediction in mice whose Tregs could not leave the tumor-draining lymph-node. Our work illustrates how mathematical modeling can guide experimentalists to perform focused experiments to understand therapeutic response, identify determinants of resistance to CBI, and refine treatment strategies.

## RESULTS

### A mechanistic mathematical model of key antitumor immunity features

We developed a mathematical model of anti-tumor and immunosuppressive mechanisms occurring within the tumor immune microenvironment. A common treatment to elicit an antitumor immune response is PD-1 inhibition. PD-1 binding to PD-L1 and PD-L2 recruits phosphatases in proximity to TCR and co-stimulatory receptors, decreasing signaling efficiency and T cell activation.^13,14^ The main target of PD-1 inhibition is CD8^+^ T cells,^15^ yet other components of the tumor immune environment, including Tregs^8,10^ and DCs,^16,17^ are directly or indirectly affected. For these reasons, we developed a mathematical model encompassing CD8^+^ T cells, Tregs, and DCs at different stages of activation. While the tumor is heterogeneous, there exist niches whereby CD8^+^ T cells, Tregs, and DCs are well-mixed with tumor cells,^10^ allowing us to specifically model mechanisms occurring within these niches using ordinary differential equations (ODEs).

To build our system of ODEs, we made assumptions rooted in current knowledge of the tumor immune environment (**Figure 1**). First, we assumed that tumor cell (N) accumulation was exponential. The tumor environment produces chemokines attracting inactive DCs (DI), tumor antigen-specific CD8^+^ effector T cells (TE), and Tregs (TR) to the tumor site. Tumor cells also present antigens, which activate CD8^+^ effector T cells, and in turn kill tumor cells. As per Breart et al., 2008, we assume that the killing capacity of CD8^+^ T cells is limited. ^18^ Upon death, tumor cells release tumor-associated antigens, which contribute to DC activation.^19^ DCs can also be activated by IFNγ produced by CD8^+^ T cells.^16^ Active DCs (DA) present antigens to both CD8^+^ T cells^20^ and Tregs,^12^ activating them. IFNγ produced by CD8^+^ T cells also upregulates PD-L1^21^ on both activated DCs and tumor cells, and the subsequent binding to PD-1 on CD8^+^ T cells results in their deactivation.^15^ In agreement with recent data, we also posited that Treg activation by inhibition of PD-1 on their surface was negligible.^410^ We factored in CD8^+^ T cell exhaustion upon repeated activation by cognate antigens presented in the tumor environment,^22^ and its reversal by IL-2.^23^ The main source of IL-2 in the tumor environment is CD8^+^ T cells,^10^ and IL-2 is modeled to be in local equilibrium, in agreement with evidence of limited diffusion of IL-2 around its producers.^24^ IL-2 is uptaken by both CD8^+^ T cells^25^ and Tregs^10^ to increase their lifespan. However, Tregs express the high-affinity IL-2 receptor at greater levels than effector T cells, outcompeting them for IL-2 usage.^26^ Treg activation inhibits CD8^+^ T cell function via adenosine production^27^ and cytokines including TGFβ, IL-10 and IL-35.^28,29^ Tregs also inhibit the expression of CD80 and CD86 on DCs via CTLA-4.^12^ These biological assumptions are translated into a system of ODEs in **Figure S1**.

**Figure 1.**
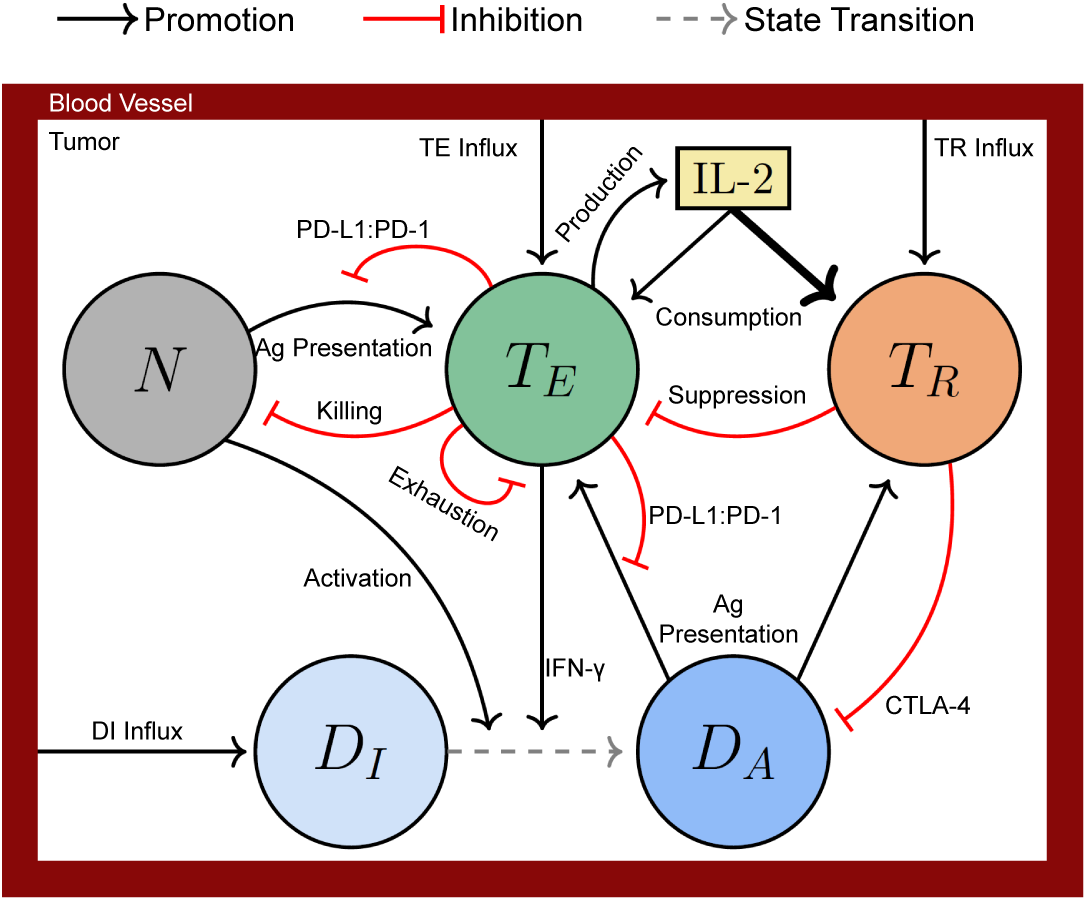
Tumor-Immune Model Schematic. Depiction of Tumor-Immune Model interactions between tumor cells (N), inactive (DI) and active (DA) dendritic cells, tumor-specific CD8^+^ T effector cells (TE), T regulatory cells (TR), and interleukin-2 (IL-2). Black arrows, promotion; red arrows, inhibition; dashed arrow, state transition. See also Figure S1.

### Mathematical model captures *in vivo* tumor growth and immune cell dynamics

We chose to apply our model to cutaneous melanoma because it is responsive to checkpoint immunotherapies, including αPD-(L)1 monotherapy, yet many patients develop acquired resistance.^2^ To enable the model to be predictive and quantitatively accurate, we parametrized it using available literature and estimated the remaining parameters by fitting newly generated experimental data. We first assessed the dynamics of tumor growth in the immunogenic melanoma cell line D4M-S, a derivative of D4M melanoma^30^ expressing the immunodominant peptide OVA_257-264_ (SIINFEKL).^31^All mice developed tumors and displayed a transient stagnation of tumor growth before progression (**Fig. 2A**). We then quantified immune cells at multiple timepoints during tumor development using flow cytometry (**Fig. 2B**). Tumor-associated DCs were recognized based on expression of MHC-II, CD11c, and CD135, and further divided into inactive or active DCs, indicated by the low or high expression of MHC molecules (**Fig. S2A**). We defined Tregs by the co-expression of CD4 and Foxp3 (**Fig. S2B**). Tumor antigen-specific CD8^+^ T cells were identified using SIINFEKL-specific tetramers. To distinguish functional from exhausted CD8^+^ T cells, we stained them with antibodies recognizing Tcf1 and Tim3, retaining cells with any expression of Tcf1 and low expression of Tim3 as functional (**Fig. S2C**). We chose such gating strategy after drawing small regions along the Tcf1 – Tim3 continuum, and quantifying concomitant production of IFNγ and TNF, a hallmark of T cell functionality,^32^ in each region (**Fig. S2D**). We set the baseline expression of IFNγ and TNF to be that of Tcf1^hi^ cells. Cytokine production peaked in Tcf1^lo^ Tim3^−^ T cells and then decreased below the baseline in Tim3^int-hi^ T cells, which we considered exhausted (**Fig. S2E**). Using these strategies, we were able to quantify how many cells populated the tumor environment during progression (**Fig. 2C**). Together with parameters found in the literature (**Table 1**), the model was fit to this original data (**Fig. 2D; Methods**), as well as to published experiments (**Fig. S2F, G**).

**Figure 2.**
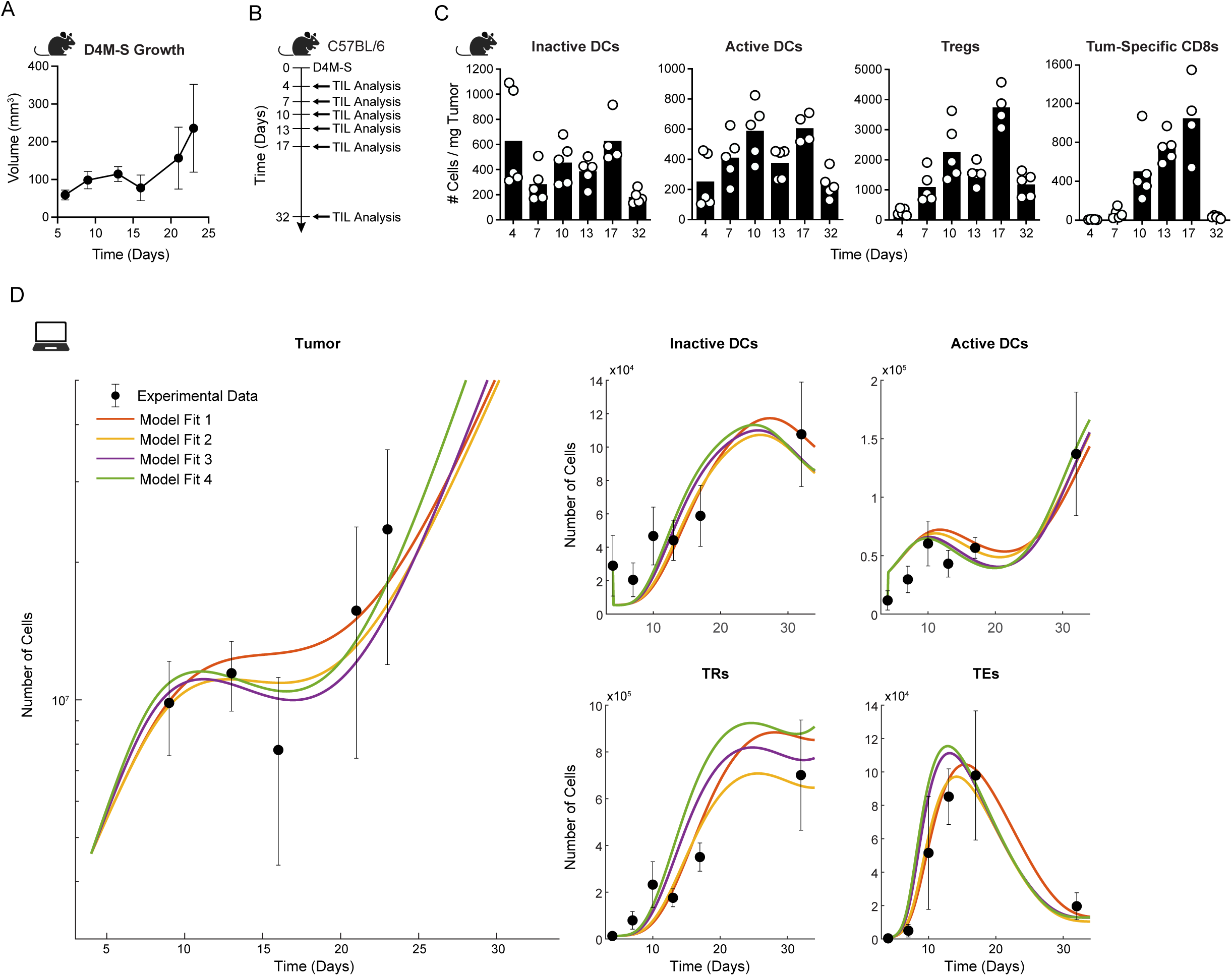
Model captures in vivo tumor growth and immune cell dynamics. (A) Mean growth curve of D4M-S tumors in mice, *n* = 5. Error bars depict one standard deviation. (B) Experimental scheme to quantify immune cell dynamics in mice bearing D4M-S melanomas. (C) Number of the indicated cell type per mg of tumor over time. *n* = 4 − 5 mice per group. Bar represents the mean number of cells. (D) Model fit to tumor (left) and immune cell (right) experimental data. Each colored line represents a unique parameter fit to the experimental data. Black dots are the mean tumor size from (A) (left) and the number of immune cells per tumor (right). Error bars depict one standard deviation. See also Figure S2.

**Table 1.**

| Parameter Values |  |  |  |
| --- | --- | --- | --- |
| Par | Description | Value/Range | Source |
| $r_N$ | Tumor proliferation rate | [0.17, 0.24] | This paper |
| $d_N$ | Tumor killing rate by $T_E$ | [12, 33] | This paper & Breart *J Clin Inv* 2008 |
| $K_N$ | Saturation of killing | 100000 | This paper |
| $\delta_I$ | $D_I$ influx rate | [0.001013, 0.004998] | This paper |
| $K_D$ | Saturation of $D_I$ influx | [1.45e-08, 8.45e-08] | This paper |
| $\sigma_R$ | $D_A$ deactivation rate by CTLA-4 on $T_R$ | 0.5688 | Hou *J Immunol* 2015; Waters *Methods Mol Biol* 2023 |
| $\rho_{DE}$ | $D_I$ activation rate via IFNγ | [0.000101, 1.483952] | This paper |
| $\rho_{DN}$ | $D_I$ activation rate via tumor antigen | [0.002504, 0.021826] | This paper |
| $K_{DN}$ | Saturation of $D_I$ activation | [4.67e-16, 5.0e-09] | This paper |
| $d_I$ | $D_I$ death rate | 0.099 | Ginhoux *J Exp Med* 2009 |
| $r_A$ | $D_A$ proliferation rate | 0.001 | This paper |
| $d_A$ | $D_A$ death rate | 0.099 | Ginhoux *J Exp Med* 2009 |
| $\mu_N$ | Tumor mutational burden | 1 | This paper |
| $\rho_E$ | $T_E$ activation rate by $D_A$ antigen presentation | [4.0e-07, 3.74e-06] | This paper |
| $K_R$ | $T_E$ suppression rate by $T_R$ | [0.0001, 0.00043] | This paper |
| $K_A$ | $D_A$ PD-L1 negative feedback rate on $T_E$ | [0.040673, 1.09984] | This paper |
| Par | Description | Value/Range | Source |
| $\rho_{EN}$ | $T_E$ activation rate by tumor presenting antigen | [5.347079, 10.9775] | This paper |
| $K_{EN}$ | Tumor PD-L1 negative feedback rate on $T_E$ | [0.011881, 1.099385] | This paper |
| $\delta_E$ | Primed $T_E$ influx rate | [0.000117, 0.019909] | This paper |
| $\delta_{EN}$ | Baseline $T_E$ influx rate | 0.0001 | This paper |
| $d_E$ | $T_E$ death rate | 0.1695 | Riberio *Proc Natl Acad Sci* 2002; Feng *Comms in Nonlin Sci and Num Sci* 2021; Burton *BioRXiv* 2023 |
| $d_{Ex}$ | $T_E$ exhaustion rate | [4.66e-09, 2.0e-08] | This paper |
| $\beta_E$ | Rate IL-2 prolongs $T_E$ lifespan | [0.001006, 0.013894] | This paper |
| $\alpha_E$ | Number of IL-2 receptors on $T_E$ | 4640 | Voisinne *Cell Reports* 2015; Hofer *Front Immunol* 2012 |
| $r_R$ | $T_R$ auto-reactivity rate | [1.03e-15, 6.56e-15] | This paper |
| $\rho_R$ | $T_R$ activation rate by $D_A$ antigen presentation | [5.0e-19, 6.22e-18] | This paper |
| $\delta_R$ | Primed $T_R$ influx rate | [0.017583, 0.089861] | This paper |
| $\delta_{RN}$ | Baseline $T_R$ influx rate | 0.0001 | This paper |
| $d_R$ | $T_R$ death rate | 0.0826 | Burton *BioRXiv* 2023; Mabarrack *Society for Leukocyte Bio* 2008 |
| $\beta_R$ | Rate IL-2 prolongs $T_R$ lifespan | [0.001006, 0.013894] | This paper |
| $\alpha_R$ | Number of IL-2 receptors on $T_R$ | 40000 | Voisinne *Cell Reports* 2015; Hofer *Front Immunol* 2012 |
| $\gamma_p$ | IL-2 production rate by $T_E$ | 864000 | Hofer *Front Immunol* 2012 |
| $\gamma_c$ | IL-2 consumption rate | 95.04 | Hofer *Front Immunol* 2012 |

To verify that our model can capture known biological behaviors, we conducted a series of *in silico* experiments using the fits from Figs 2D and S2F-G. First, we simulated decreased immunogenicity and found that tumors grew exponentially, which matched the behavior seen in non-immunogenic D4M tumors (**Fig. 3A**). The unrestricted growth of D4M-S tumors upon depletion of CD8^+^ T cells in vivo was also successfully simulated using our model (**Fig. 3B**). The model also captures the expected tumor rejection upon depletion of Tregs (**Fig. S3A**).^33^ A distinguishing feature of our model is that it can capture immunosuppressive reactions triggered by checkpoint immunotherapies, such as the tumor Treg increase that follows treatment with αPD-1 (**Fig. 3C**) or αCTLA-4 (**Fig. S3B**) in both mice and patients.^10–12^ Having verified that our model can recapitulate previously described immune reactions, we used Latin Hypercube sampling^34^ to generate hundreds of parameter sets, each representing a unique virtual mouse, that fit the data equally as well to capture variability across individuals and to use for further simulations and analyses (**Fig. S3C**).

**Figure 3.**
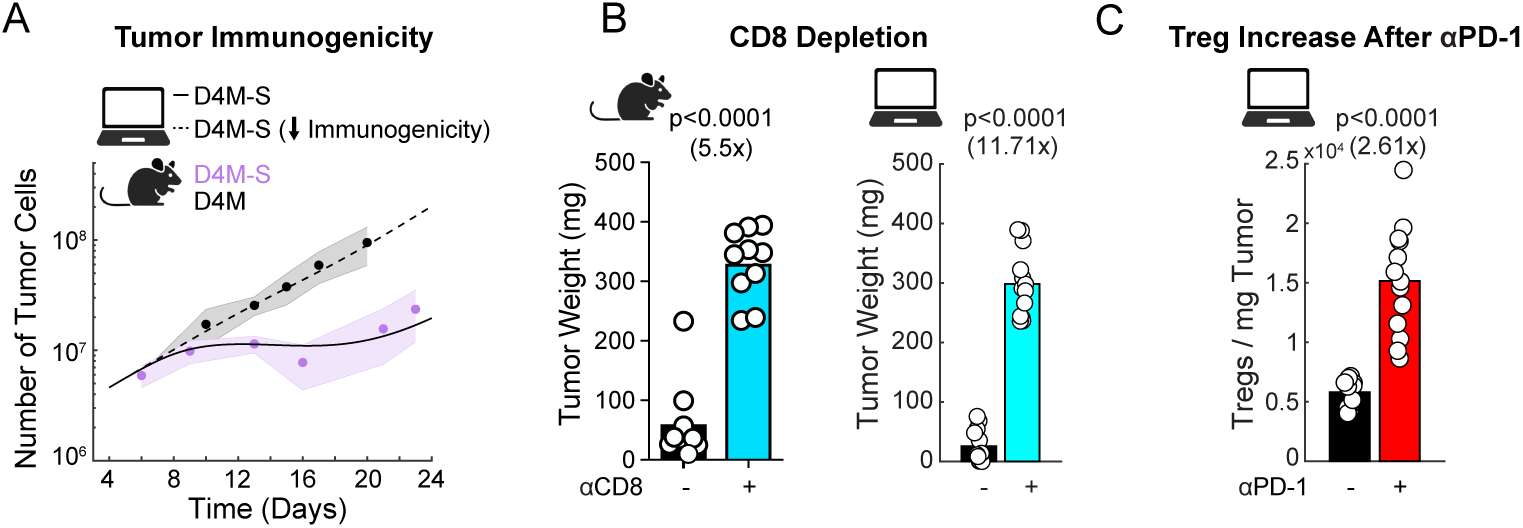
Model recapitulates known biological behaviors. (A) Unaltered model simulations (solid line, mean of fits from Fig 2D) fit to immunogenic D4M-S tumor growth data from Fig 2A (purple dots) compared to model simulations with decreased immunogenicity (dashed line), which is now comparable to non-immunogenic D4M tumor growth data (black dots). D4M tumor growth data *n* = 4 mice. Clouds around experimental data depict one standard deviation. (B) Tumor weight in mice with D4M-S melanomas treated or not with FTY720 and CD8-depleting mAbs (left). *n* = 9 − 10 mice per group. Model simulations (*n* = 13/group) are depicted on the right. (C) Simulated number of Tregs per mg of tumor, in the presence or absence of *α*PD-1 mAbs. *n* = 13/group. Compare to Figure S7N of Geels et al, Cancer Cell 2024. In B and C bars depict mean, *p* values determined by Mann-Whitney U test. See also Figure S3.

### Differential impact of tumor and immune initial conditions on tumor growth dynamics

In D4M-S tumor growth experiments we typically observe transient stagnation of growth before progression. However, in a minority of the experiments (2/7) the tumor grew monotonically (**Fig 4A**, experimental data). To determine the cause of varying D4M-S tumor growth behaviors, we investigated how early tumor immune interactions impact tumor development by perturbing the initial conditions of the tumor and immune cells. When we fixed the parameter values and increased the initial conditions of all cell types, we found that the tumors exhibited monotonic growth, suggesting that the variability of D4M-S growth behavior may be due to the initial conditions of the tumor and immune system (**Fig. 4A**, simulations). This result held true if we increased the tumor initial condition alone (**Fig. S4A**). Thus, an increased initial tumor size is sufficient for the tumor to escape immune regulation.

**Figure 4.**
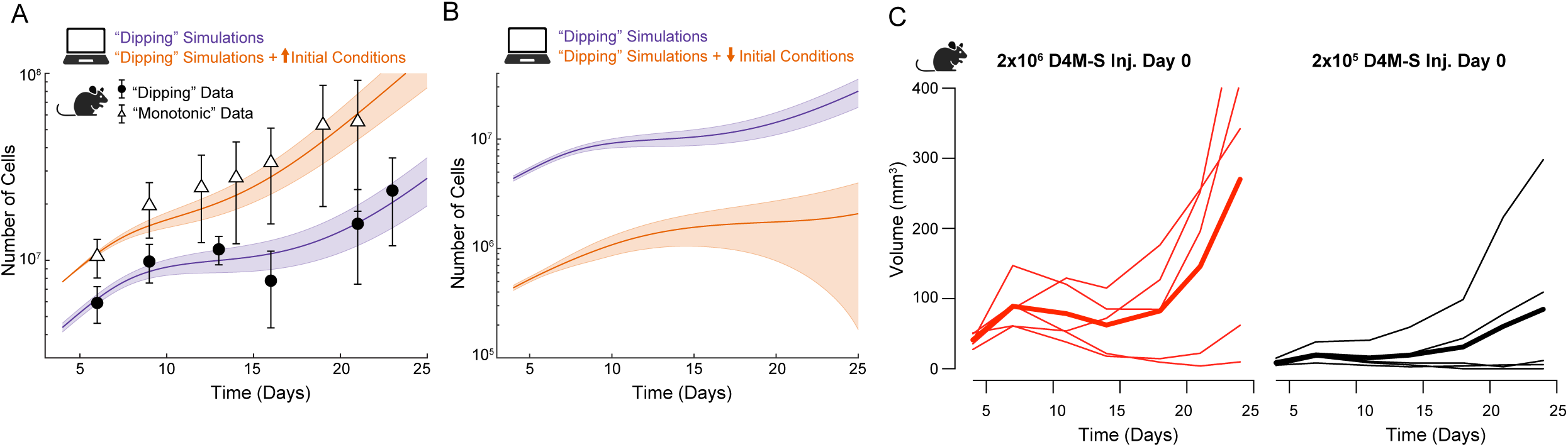
Initial tumor and immune conditions contribute to tumor outcome. (A) Unaltered model simulations (purple line, mean of fits from Fig S3C) fit to D4M-S “dipping” tumor growth data from Fig 2A (black circles). Model simulations with increased tumor and immune cell initial conditions (orange line), are comparable to D4M-S “monotonic” tumor growth data (white triangles). D4M-S “monotonic” tumor growth *n* = 5 mice. Clouds around mean simulations and error bars on data depict one standard deviation. (B) Unaltered model simulations (purple line, mean of fits from Fig 2D) and model simulations with decreased tumor and immune cell initial conditions (orange line). Clouds around mean simulations depict one standard deviation. (C) Growth curves of D4M-S tumors in mice injected with 2×10^6^ (left, red) and 2×10^5^ (right, black) tumor cells. Thin lines, individual mice; thick lines, mean of group; *n* = 5 mice/group. See also Figure S4.

We also explored the alternative hypothesis that tumor progression could occur given changes in parameter values. To this end, we fit our model parameters to data that displayed monotonic tumor growth (**Fig. S4B**). However, applying the initial conditions of the stagnated tumor growth to these parameters did not result in tumor stagnation (**Fig. S4C**), suggesting that the parameterization of the monotonic growth data does not capture all the tumor growth behaviors we observed experimentally. This further indicated to us that parameterization based on the stagnated tumor growth was more general, in that combinations with different initial conditions could capture all the observed tumor growth patterns. To corroborate this notion, we simulated the growth dynamics of D4M-S tumors upon decreasing the initial conditions of each cell type, finding decreased tumor growth (**Fig. 4B**). We validated these predictions experimentally (**Fig. 4C**), further indicating that size of the tumor and the quantity of immune cells play a role in D4M-S growth dynamics.

We next sought to understand how perturbations to the immune cell initial conditions alone could impact tumor outcomes. The infiltration of immune cells into incipient tumors is a key determinant of their outcome. For instance, early infiltration by Tregs instructs tumor growth, while immediate infiltration of CD8^+^ T cells, facilitated by vaccination, leads to cancer rejection.^35^ It is conceivable that immune interventions such as the adoptive transfer of genetically modified (i.e., CAR-T cells) or antigen-specific T cells alter the immune environment of nascent tumor masses. Thus, we varied the initial conditions of each immune cell type, both individually and in combination, and quantified the mean tumor size of all simulations on day 35 (**Fig. 5A and S5**). Interestingly, this analysis showed that small increases in the initial T effector cell population did little to decrease the mean tumor size. To see a substantial reduction in tumor, the initial T effector cell number needed to be increased at least six-fold. Conversely, small decreases to the initial conditions of Tregs, or increases in inactive and active DCs, effectively restricted tumor growth. When looking at combinations of initial immune cell alterations, there exists a threshold in which increasing the T effector cell initial condition while also increasing the Treg initial condition results in a worse outcome, even when the T effector cell initial conditions are increased more than that of the Tregs; it is not until after increasing the T effector cell initial condition past this threshold that the model predicts improved tumor outcomes (**Figure S5**). The model further suggests that the DCs modulate much of the immune response, as any combination of increased initial conditions of inactive and active DCs results in decreased tumor size.

**Figure 5.**
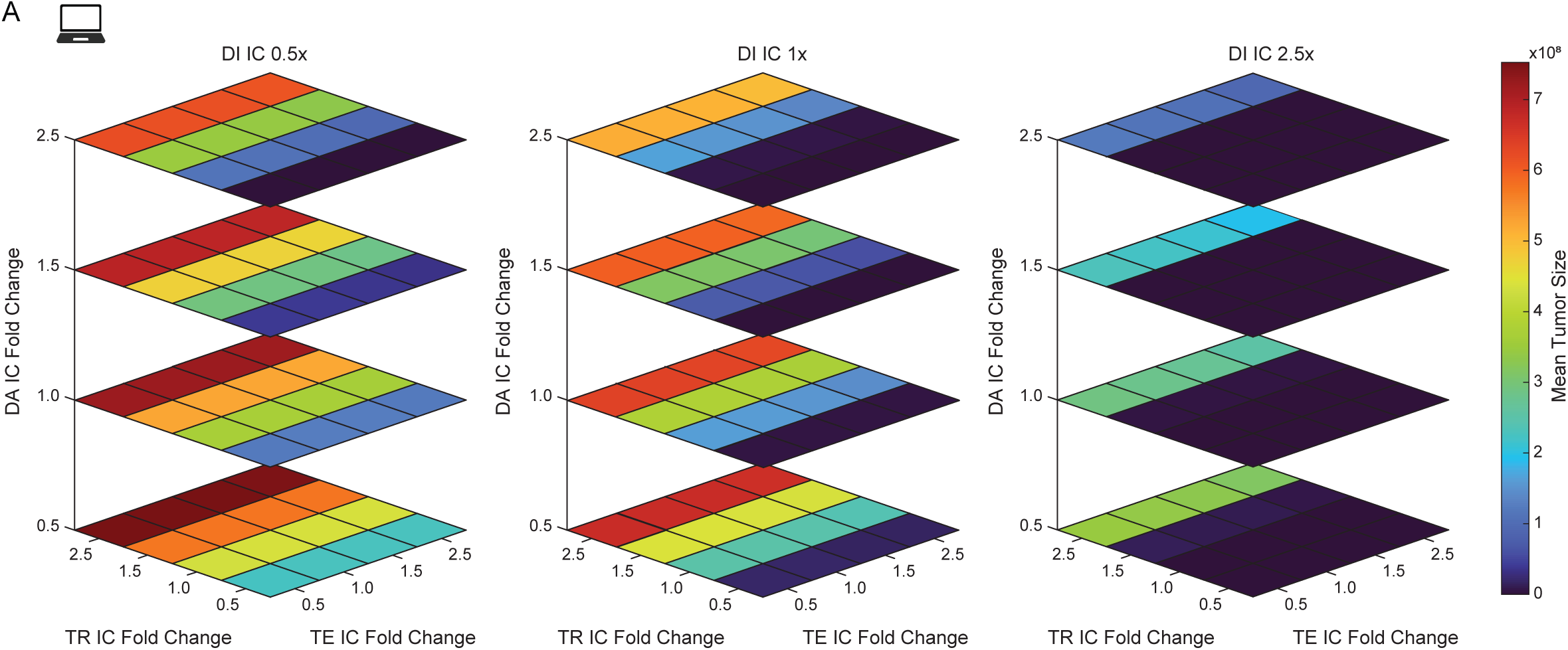
Specific alterations in immune cell initial conditions result in increased tumor control. (A) Mean tumor size at final simulation time (day 35) for the indicated fold change in TE, TR, DA, and/or DI initial condition. *n* = 299 simulations/square. See also Figure S5.

### Mathematical model accurately simulates the experimental response to *α*PD-1

We next investigated whether our mathematical model could recapitulate the experimental response to PD-1 immunotherapy. We reanalyzed the growth curves of αPD-1-treated D4M-S tumors (published in ^10^), and classified mice as complete responders if they fully rejected the tumor and non-responders otherwise (**Fig. S6A**). Then we mathematically simulated *⍺*PD-1 immunotherapy by reducing the values of the parameters K_A_ and K_EN_, representing the deactivation of CD8^+^ T cells by PD-L1 expressed from active DCs and tumor cells (**Fig. S1**). We found that 25% PD-1 blockade allowed tumor rejection in a minority of the virtual mice, while 75% blockade led to tumor rejection in almost all cases (**Fig. S6B-C**). Notably, a PD-1 blocking efficiency of 47% successfully allowed the model to recapitulate the 30% responder / 70% non-responder ratio seen in the experimental data (**Fig. 6 and S6D**). Thus, our mathematical model accurately recapitulated the outcome of *⍺*PD-1 immunotherapy in mice. Moreover, our simulations indicated that an increasingly efficient PD-1 blockade would correlate with enhanced antitumor efficacy.

**Figure 6.**
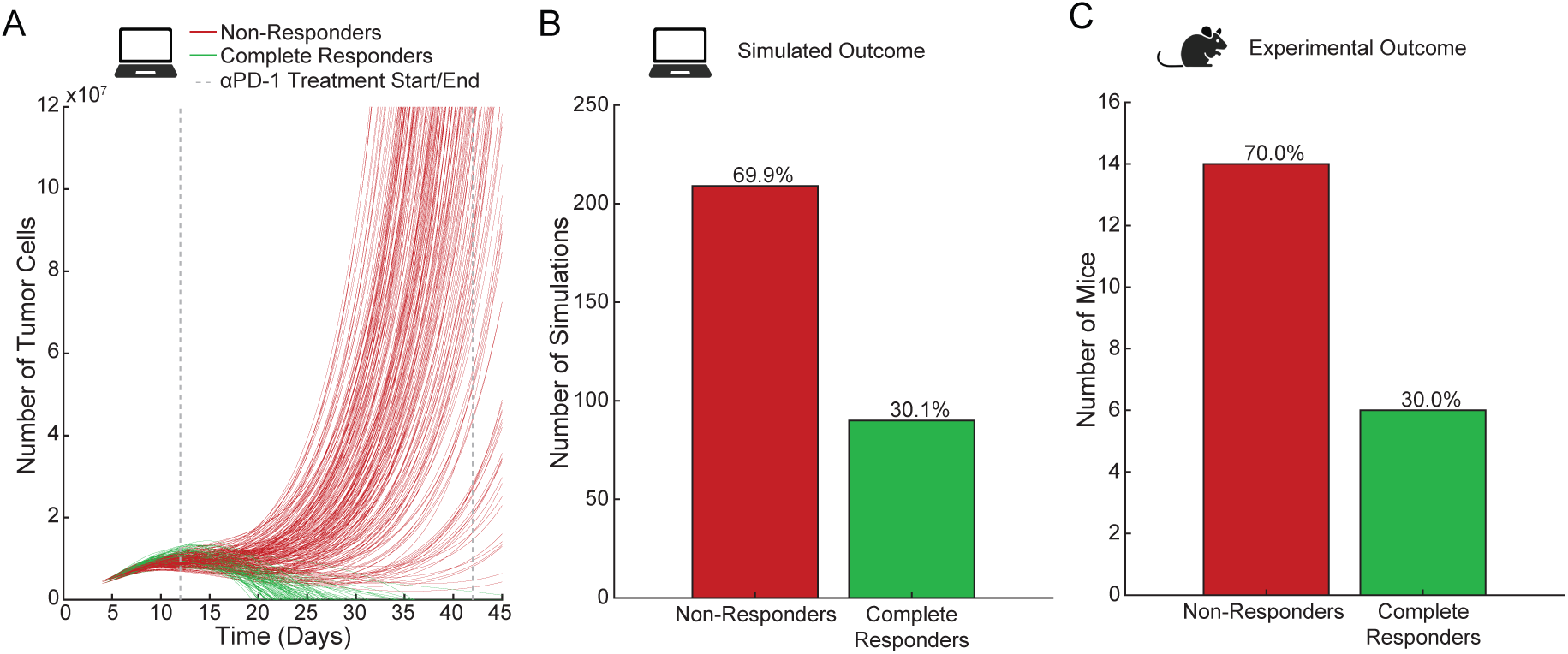
Model captures the experimental response to *α*PD-1. (A) Simulated response to *⍺*PD-1 treatment classified as “Complete Responders” (green, tumors that are fully rejected) and “Non-Responders” (red, all other tumors). Dashed lines indicate treatment start and stop time. *⍺*PD-1 blocking efficiency is set to 47%; *n* = 299 simulations. (B) Simulated outcome to *⍺*PD-1 treatment by response type. (C) Experimental outcome to *⍺*PD-1 treatment by response type, as displayed in Figure S6A. See also Figure S6.

### Treg influx hinders response to *α*PD-1

To identify key determinants of acquired resistance to PD-1 immunotherapy, we used our *⍺*PD-1-treated simulations and first characterized the differences between responders and non-responders by identifying which parameters took significantly different values in responders and non-responders (**Fig. 7A and S7**). Only five parameters showed statistically significant differences between the two groups: *δ*_*R*_ (Treg influx rate), *r*_*N*_ (tumor growth rate), *r*_*E*_ (antigen presentation from active DC to CD8^+^ T cells), *d*_*E*_ (influx of primed CD8^+^ T cell), and *d*_*Ex*_(degree of CD8+ T cell exhaustion) (**Fig. 7A**). Next, to identify additional interventions that would enhance the therapeutic efficacy non-responders, we decreased each parameter individually by 5% and quantified the total change in response to *⍺*PD-1 immunotherapy as compared to the original simulations (**Fig. 7B**). We found that four individual parameters greatly increased the response rate, namely *K*_*A*_ (PD-L1 amount in active DCs), *K*_*R*_ (direct suppression of CD8^+^ T cell function by Tregs), *δ*_*R*_, and *r*_*N*_. Since *K*_*A*_ and *K*_*R*_ were not statistically significant when we assessed the differences in responder and non-responder simulations (**Fig. S7**), we excluded them from further analysis. We also excluded *r*_*N*_, the tumor proliferation rate, from further analysis as decreasing its value and seeing improved tumor control is trivial. Thus, the only remaining parameter that was both statistically different between complete and non-responders and greatly increased the response of non-responders to *⍺*PD-1 immunotherapy when decreased was *δ*_*R*_, the Treg influx rate. To experimentally validate that a reduction in the Treg influx to the tumor could enhance the response to *⍺*PD-1 immunotherapy, we generated mice in which approximately half of the Tregs are deficient in S1PR1, a receptor necessary for leaving the tumor-draining lymph nodes.^10^ To do so, we irradiated CD45.1 mice and reconstituted them with a 1:1 mixture of Foxp3^DTR^ and either *S1pr1*^f/f^ (CTRL) or *S1pr1*^f/f^ x UBC^creERT2^ (S1PR1^KO^) bone marrow. Treatment with tamoxifen would delete S1PR1 from half of the lymphocytes, preventing their egress from the tumor-draining lymph node to invade the tumor. At the same time, diphtheria toxin (DT) administration would deplete normal Tregs, but not other cells, making S1PR1 deficiency specifically targeted at Tregs. Upon hematopoietic reconstitution, we challenged mice with D4M-S tumors, administered αPD-1 or an isotype control starting on day 12, and followed the dynamics of tumor growth (**Fig. 7C**). We analyzed the change in tumor size after the start of immunotherapy of each experimental group. In general, isotype control antibodies did not prevent tumor progression in both CTRL and S1PR1^KO^ mice, even if one mouse experienced delayed progression and one showed tumor rejection in the S1PR1^KO^ group (**Fig. 7D**). Administration of PD-1 immunotherapy to control animals resulted in transient tumor control, yet progression ensued in most of the mice. Conversely, PD-1 inhibition in the S1PR1^KO^ group was able to eliminate or significantly dampen tumor growth (**Fig. 7D**) and enhance mouse survival (**Fig. 7E**). Therefore, our mathematical model identified the most promising aspect of tumor immunity (out of 33 parameters) that synergized with PD-1 blockade to delay or avoid the development of secondary resistance to αPD-1 immunotherapy.

**Figure 7.**
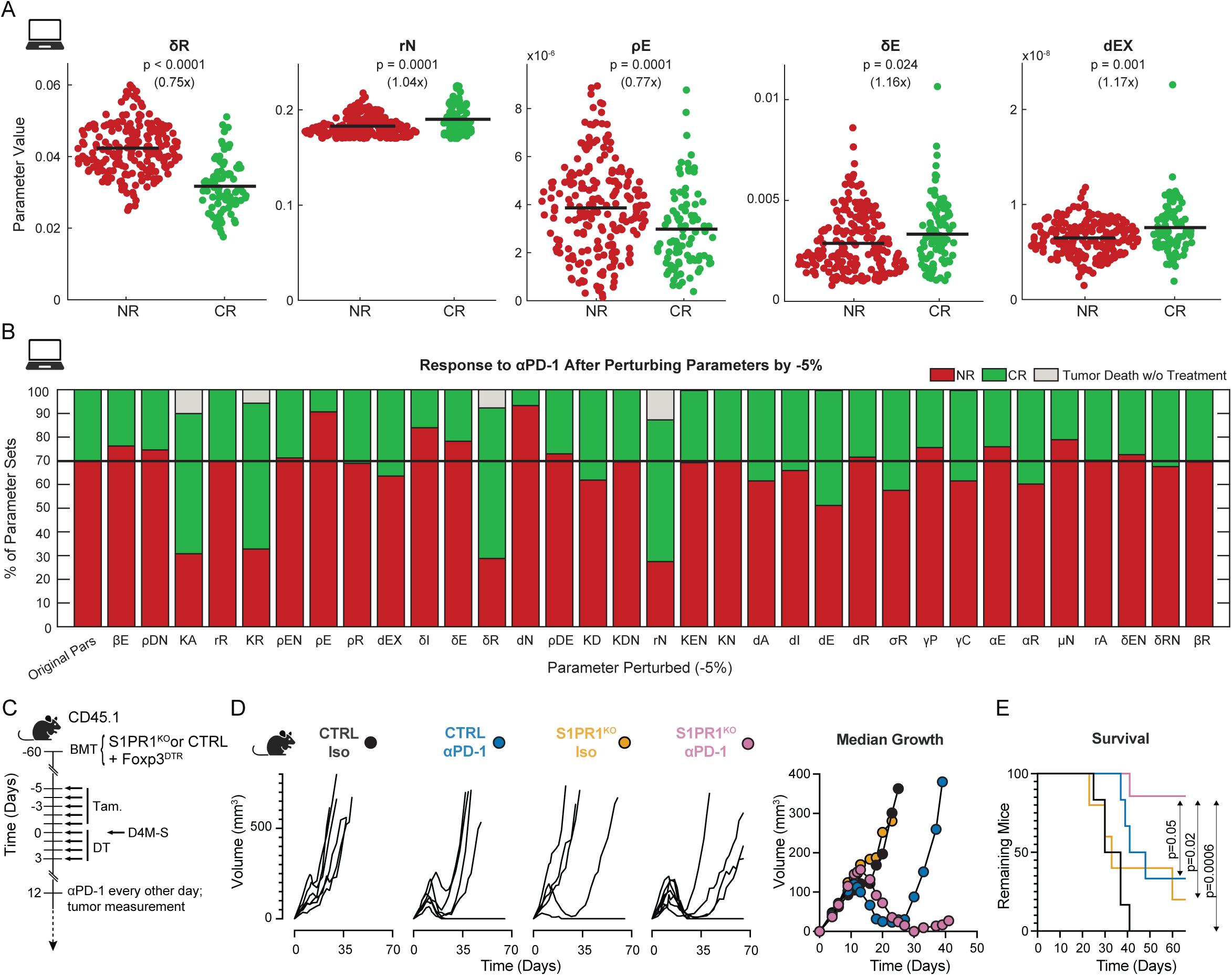
Large Treg influx hinders response to *α*PD-1. (A) Difference between parameter values in simulations classified as “Non-Responders” (red) and “Complete Responders” (green) with simulated *⍺*PD-1 treatment. Black bars depict mean; *n* = 299 simulations; *p* values determined by Mann-Whitney U test. (B) Simulated response to *⍺*PD-1 treatment with blocking efficiency set to 47% after decreasing an individual parameter by 5% of its original value. “Non-Responders”, red; “Complete Responders”, green; simulations that resulted in tumor death by only decreasing parameter value, gray. *n* = 299 simulations/bar. (C) Experimental scheme for Treg influx depletion. (D) D4M-S tumor growth in mice with or without decreased Treg influx rates in the presence or absence of *⍺*PD-1. Individual mice shown in each group (left) and median tumor growth on the last day all mice within a group were alive (right). *n* = 6 − 7 per group. (E) Survival curves of the mice in D. *n* = 6 − 7 per group. P values by Mantel-Cox test. See also Figure S7.

## DISCUSSION

In this study, we developed a mechanistic mathematical model grounded in current biological knowledge to identify the most critical immunological features that influence tumor growth. Once we successfully fit the model to experimental data, we ensured that it could recapitulate known biological behaviors such as increased tumor growth in non-immunogenic cancers and upon depletion of T effector cells, increased tumor control upon depletion of Tregs, and the immunosuppressive reactions triggered by checkpoint immunotherapies. By conducting simulations on hundreds of ‘virtual mice’ to capture biological variability, we identified the engraftment of tumor cells and specific alterations to the immune system early after tumor establishment as critical factors influencing tumor control or escape. Crucially, we used the model to simulate *⍺*PD-1 immunotherapy and identified Treg influx as a key determinant of acquired resistance to it. Our *in silico* predictions were validated using focused experiments.

Several mathematical models of the tumor immune system have been previously developed and have contributed to the investigation of mechanisms of cancer control.^36–40^ Some of these models include immunomodulatory mechanisms,^41–44^ yet none consider how checkpoint immunotherapy supports immunosuppression. Distinctive features of our mathematical model are that it considers how antigen-presenting cells^12^ and effector T cells^10^ support tumor Tregs, the effect of IL-2 on T cell exhaustion,^23^ and how these aspects are influenced by checkpoint blockade. These factors allowed us to capture the Treg increase observed after checkpoint immunotherapy,^10,12^ which is key to identifying the determinants of acquired resistance to CBI. As is the case with all mathematical models, we did not include every possible cell type in the tumor environment or all mechanisms of interaction between them. This leaves ample room for model expansion to consider additional immunosuppressive effects, such as macrophage polarization away from pro-inflammatory functions,^45^ chronic IFN signaling,^46,47^ or the accumulation of T regulatory type 1 (Tr1) cells.^48^

Since ODE simulations typically model a variable over time, it is useful to have temporal data to fit the model to and enable quantitative analysis and predictions. However, time-course experiments can quickly become expensive and time-consuming. We conducted a pseudo-temporal experiment to quantify immune cells across tumor development by sacrificing five mice at each time point of interest (**Fig. 2B and C**). This experiment provided a correlation between time and immune cell counts but could not examine cell counts longitudinally within the same animal. Such repeated-measures experiments are typically limited to monitoring tumor growth or collecting samples from accessible tissues (e.g., blood) over time. Recently, intravital dynamic microscopy emerged as a technology providing time-and space-resolution of cell motility^49,50^ and activation,^10,12,51^ and allowed longitudinal imaging of the same area days apart.^12,52^ We foresee that intravital microscopy will become a prime source for spatio-temporal data for modeling purposes in the future.

Our model shows that the number of tumor and immune cells at early time points (‘initial conditions’) predicts cancer control or escape (**Figs. 4 and 5**). We speculate that this observation can be extended to changes in the tumor immune environment before or immediately after a major immunological intervention (e.g., CBI). In support of this statement, a recent report demonstrated that recent COVID-19 vaccination improved the outcome of CBI.^53^ While type I IFN production had a major role, it would be interesting to test whether mRNA vaccines alter ‘initial conditions’ to sensitize cancers to checkpoint immunotherapy. Moreover, it has been described that early-on-therapy changes in tumor size^54^ or the composition of the tumor immune infiltrate^55^ predict response to immunotherapy. Thus, how could ‘initial conditions’ be manipulated to maximize the effectiveness of CBI? In a perspective article, Dr. Lieping Chen argued that increasing immune effector function in the presence of immunosuppressive mechanisms may not result in tumor control, akin to increasing pressure in a clogged water pipe.^56^ Accordingly, our model and the work of another group^57^ predict that the inhibition of immunosuppressive cells leads to better tumor control than potentiating effector cells. This notion finds experimental correlates in the increased effectiveness of PD-1 blockade after immune interventions to limit Treg response, such as inhibition of ICOSL,^10^ administration of depleting αCTLA-4 mAbs,^58^ or treatment with cyclophosphamide.^59^

We predict that an increasingly efficient PD-1 blockade would correlate with enhanced antitumor efficacy (**Fig. S6**). One of the most commonly used αPD-1 mAbs in humans, nivolumab, occupies PD-1 at 60-80% efficiency at the clinically relevant doses of 0.3 to 3 mg/kg,^60^ while computer simulations predict pembrolizumab to be slightly more efficient.^61^ Thus, there is an unmet need and a compelling rationale to generate αPD-1 mAbs with enhanced affinity for clinical use. Notably, the recently developed antibody, sintilimab, has greater affinity for PD-1 than nivolumab and pembrolizumab.^62^ The therapeutic efficacy of these mAbs is currently being compared, with initial results in NSCLC patients indicating at least equivalent performance.^63^

The comparison of parameter values between virtual mice responding or not to PD-1 blockade (**Fig. 7A**) and our parameter perturbation analysis (**Fig. 7B**) converged on Treg influx into the tumor as a key determinant of resistance to PD-1 CBI. We validated this prediction experimentally (**Fig. 7C, D**), providing a compelling rationale for developing drugs that block Treg recruitment to tumors. Unfortunately, this task is difficult due to the lack of specificity and the redundancy of the chemokine system. Antibodies targeting CCR4, necessary for Treg homing to a variety of cancers,^64^ did not synergize with PD-1 CBI because they inhibited both Tregs and antitumor CD8^+^ T cells.^65^ Moreover, CCR8, expressed specifically in tumor Tregs, may not be necessary for Treg homing to tumors.^66^ These limitations might be overcome by the development of bispecific antibodies capable of targeting activated Tregs (e.g., αCCR8, αOX-40, αCTLA-4) while blocking chemokine receptors guiding Tregs into cancers, such as CCR4, CCR5, or CXCR3.^64^ Our simulations also provided other valuable insights into the mechanisms of PD-1 immunotherapy against immunogenic melanoma. Since increased tumor cell death (*d*_*N*_) and tumor antigen presentation by DCs (*ρ*_*E*_) correlate with better melanoma control, we speculate that PD-1 blockade will synergize with immunogenic chemotherapy, as shown in lung cancer models.^67^ Additionally, PD-L1 may be more immunosuppressive when expressed by DCs (*K*_*A*) than tumor cells (*K*_*EN*_), as observed in a mouse model of colon cancer.^68^ Finally, our analysis sheds some light on the relative importance of Treg-mediated inhibition of APC (*σ*_*R*_) or effector T cells (*K*_*R*_) during PD-1 CBI: the latter may play a more prominent immunosuppressive role. This hypothesis awaits experimental validation.

Recent studies have shown 10-year progression-free survival in 31% of metastatic melanoma patients treated with combination *⍺*PD-1 and *⍺*CTLA-4 as compared to 23% with *⍺*PD-1 or 6% with *⍺*CTLA-4 alone.^1^ While this is a major improvement from previous metastatic melanoma treatments, the majority of patients eventually cease to respond to state-of-the-art checkpoint blockade immunotherapy. This consideration illustrates the need for further studies to uncover optimal therapeutic strategies. Seminal work showed that it is possible to use mathematical modeling to predict the optimal temporal combinations of preventive antitumor vaccination in mouse models.^69^ While our current work focuses on *⍺*PD-1 monotherapy, the model we present here can simulate CTLA-4 inhibition, blockade of effector T cell help to Tregs (similar to the effect of *⍺*ICOSL^10^), or restriction of T cell exhaustion (resembling a possible mechanism of action of *⍺*LAG-3^22^), and can be used to identify combinations for optimal therapeutic efficacy in a time-efficient manner.

## ACKNOWLEGEMENTS

This work is funded by pilot grants from the Chao Family Comprehensive Cancer Center (CFCCC) and the Cancer Systems Biology Center of the University of California Irvine, and the DoD Team grant ME220176P1 (F.M.). R.S.S. and S.N.G. are supported by NSF-GRFP fellowships 1000330047 and 2235784, and A.M. by the NIH-NIAMS F31 AR083279. S.A.V. is supported by R01 NS120060, which also provided a diversity supplement to R.S.S. J.S.L wishes to acknowledge support from DMS-1763272 and the Simons Foundation (594598QN) for an NSF-Simons Center for Multiscale Cell Fate Research, as well as grants P30 CA062203 and P01CA288662.

## CONTRIBUTIONS

R.S.S. devised and analyzed the mathematical model and wrote the paper. S.N.G, A.M, and C.M. performed wet-lab experiments. S.A.V. provided scientific guidance. J.S.L. and F.M supervised the study and wrote the paper.

---

**Figure S1.**
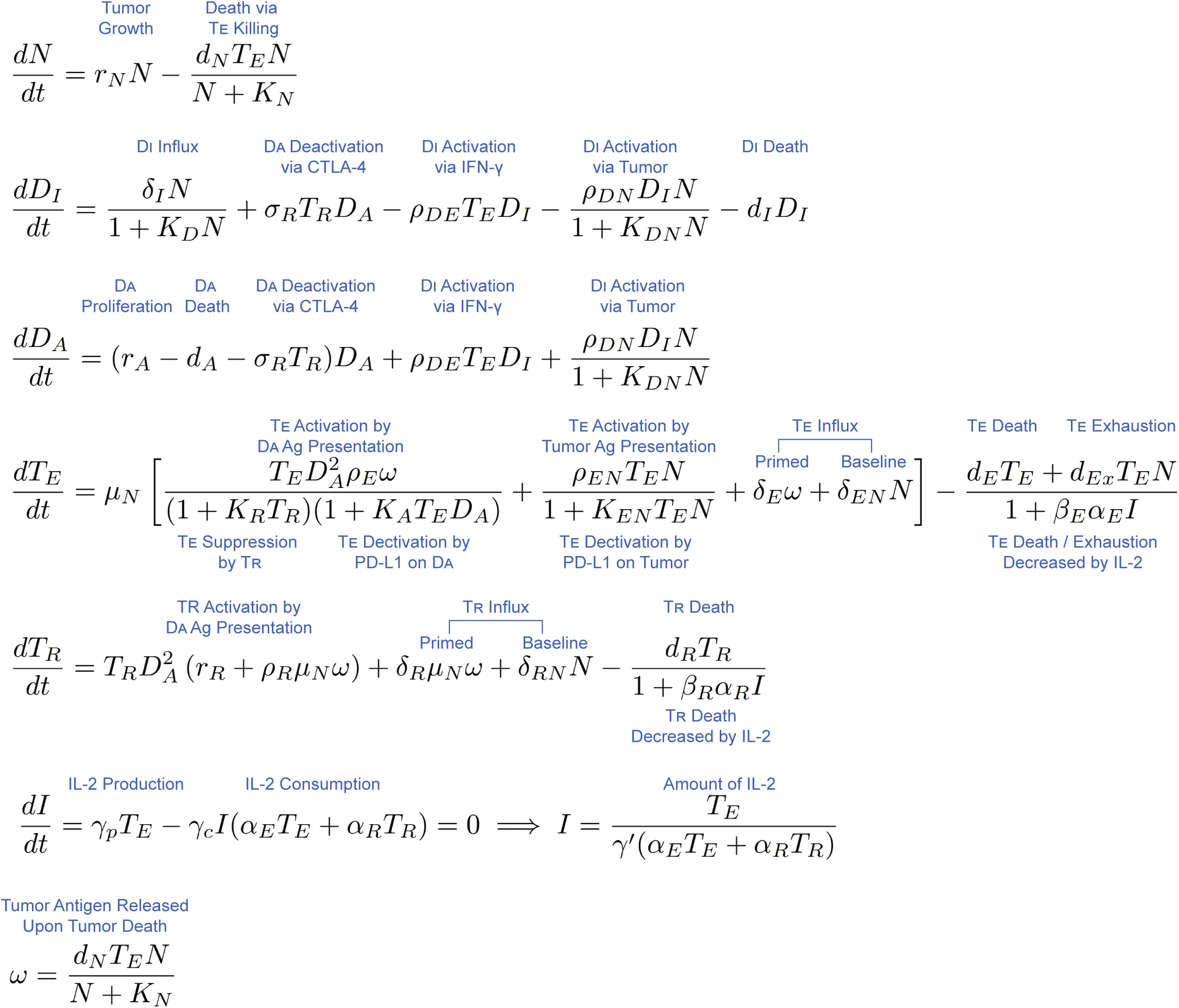
Ordinary differential equations for the tumor-immune model. Model equations with annotation describing the mechanism represented by each term.

**Figure S2.**
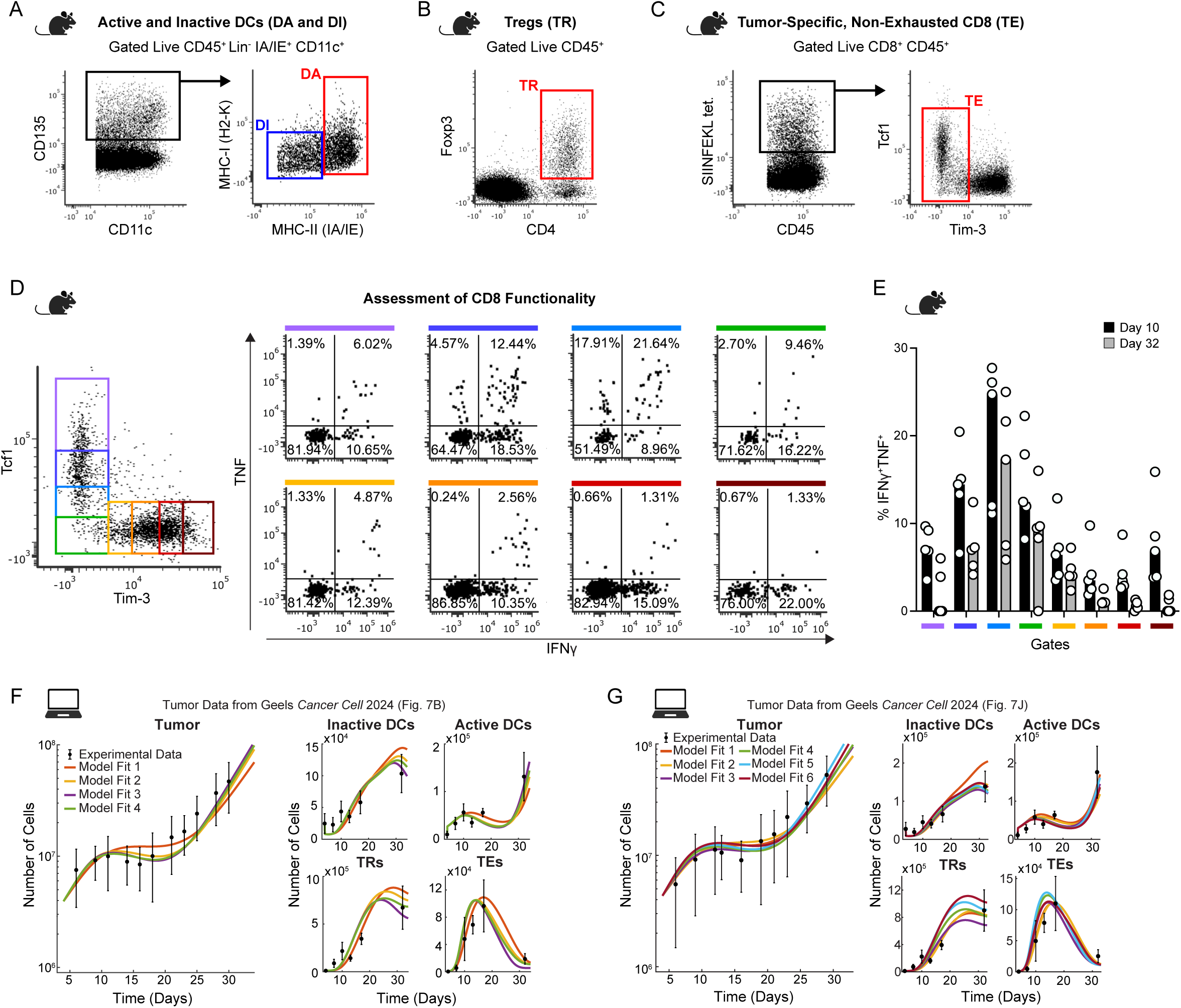
Model captures previously published in vivo tumor growth and immune cell dynamics, related to Figure 2. (A) Flow cytometry gating to identify active (DA) and inactive (DI) dendritic cells. (B) Flow cytometry gating to identify T regulatory cells (TR). (C) Flow cytometry gating to identify non-exhausted CD8+ T cells (TE), based on the results from D and E. (D) Flow cytometry gating to identify non-exhausted CD8+ T cells. Discreet gates along the TCF-1 / Tim 3 continuum (left) were applied to assess the functionality of CD8+ T cells. The readout of functionality was the production of IFNγ and TNF after in vitro restimulation of tumor-infiltrating lymphocytes with αCD3 and αCD28 (right). Gates are color-coded. (E) Summary of CD8+ T cells producing both IFNγ and TNF 10 and 32 days after tumor injection. *n* = 5 mice per group. Bars represent the mean. (F and G) Model fit to tumor growth data from Geels *Cancer Cell* 2024 (left) and immune cell experimental data (right). Each colored line represents a unique parameter fit to the experimental data. Black dots are the mean tumor size from two experiments with *n* = 4 − 5 / experiment (left) and the calculated number of immune cells per tumor (right). Error bars depict one standard deviation.

**Figure S3.**
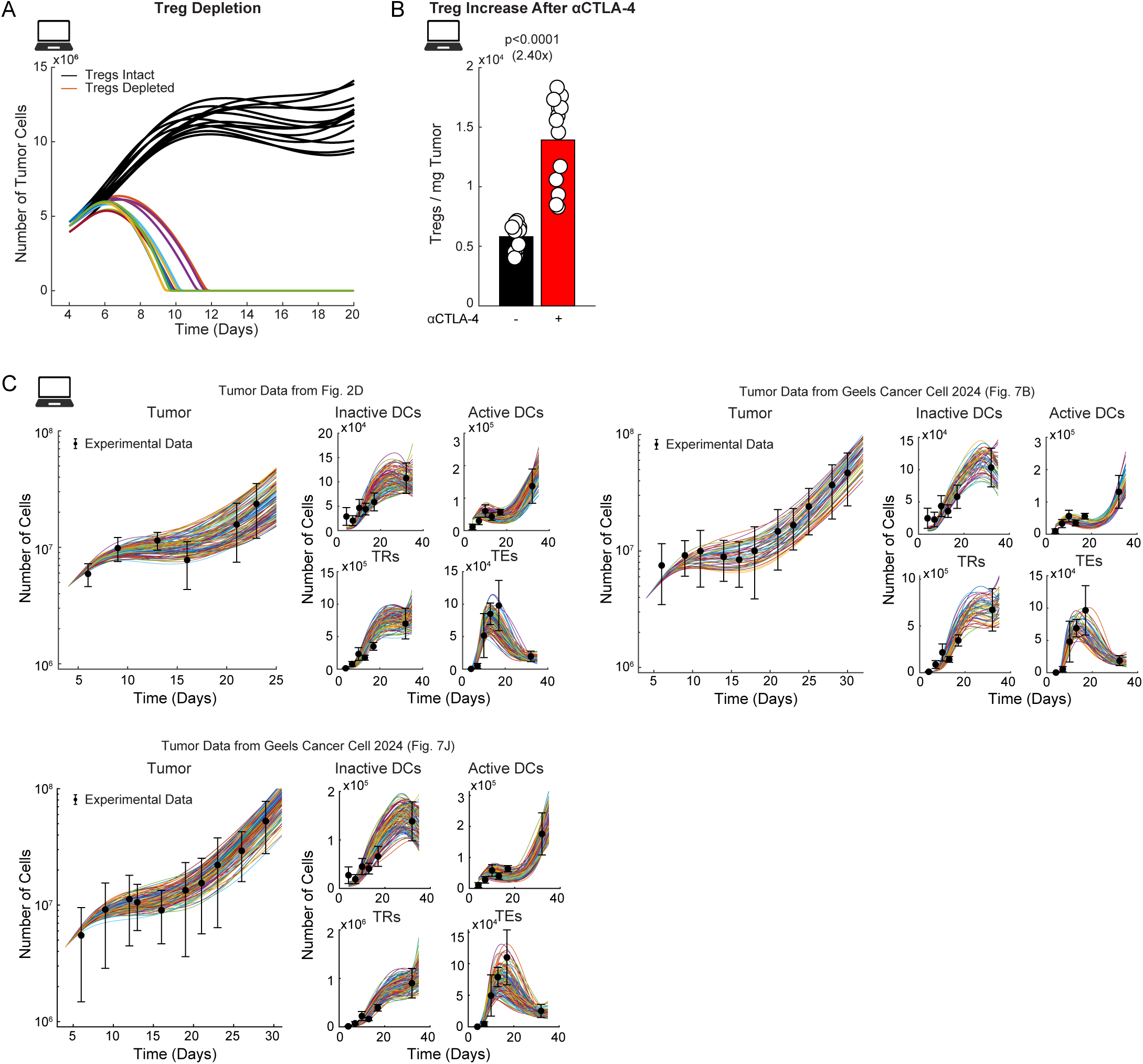
Generation of fits using Latin Hypercube Sampling, related to Figure 3. (A) Simulation of tumor growth in presence (black lines) or absence of Tregs (colored lines). Model simulations *n* = 13/group. (B) Simulated number Tregs per mg of tumor in the presence or absence of CTLA-4 blockade. Bars depict mean, *p* values determined by Mann-Whitney U test, and model simulations *n* = 13/group. (C) Parameter sets generated via Latin-Hypercube Sampling, shown with experimental data. 117 LHS parameter sets that fit experimental data from Figure 2D (top left), 53, and 129 parameter sets that fit experimental data from Geels Cancer Cell 2024 (top right and bottom panels are based on data in Fig. 7B and 7J, respectively).

**Figure S4.**
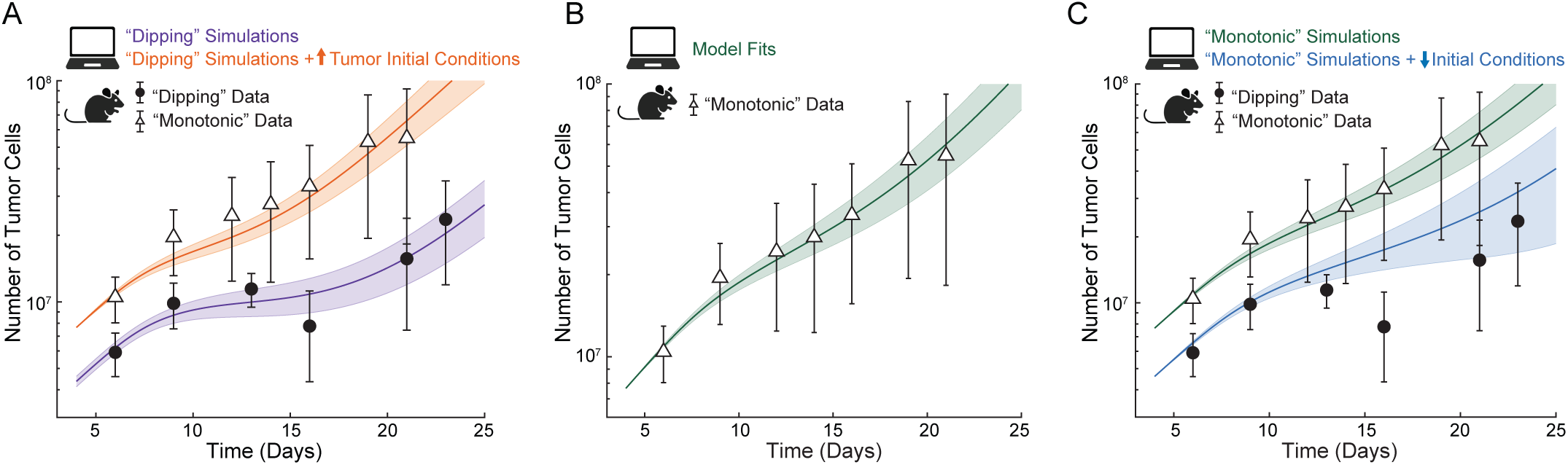
Model predictions after changing the initial conditions for tumor cell numbers only, and fit to monotonic tumor growth data, related to Figure 4. (A) Unaltered model simulations (purple line, mean of fits from Fig 2D) fit to D4M-S “dipping” tumor growth data from Fig 2A (black circles) and model simulations with increased tumor initial conditions (orange line), which is now comparable to D4M-S “monotonic” tumor growth data (white triangles). D4M-S “monotonic” tumor growth data *n* = 5 mice. Clouds around mean simulations and error bars on data depict one standard deviation. (B) Mean of model fits (green line) to “monotonic” D4M-S tumor growth data (white triangles). Model simulations *n* = 618; D4M-S “monotonic” tumor growth data *n* = 5 mice. Clouds around mean simulations and error bars on data depict one standard deviation. (C) Unaltered “monotonic” model simulations (green line) fit to D4M-S “monotonic” tumor growth data (white triangles). Model simulations with decreased tumor and immune cell initial conditions (blue line), which are not comparable to D4M-S “dipping” tumor growth data (black circle). Clouds around mean simulations and error bars on data depict one standard deviation.

**Figure S5.**
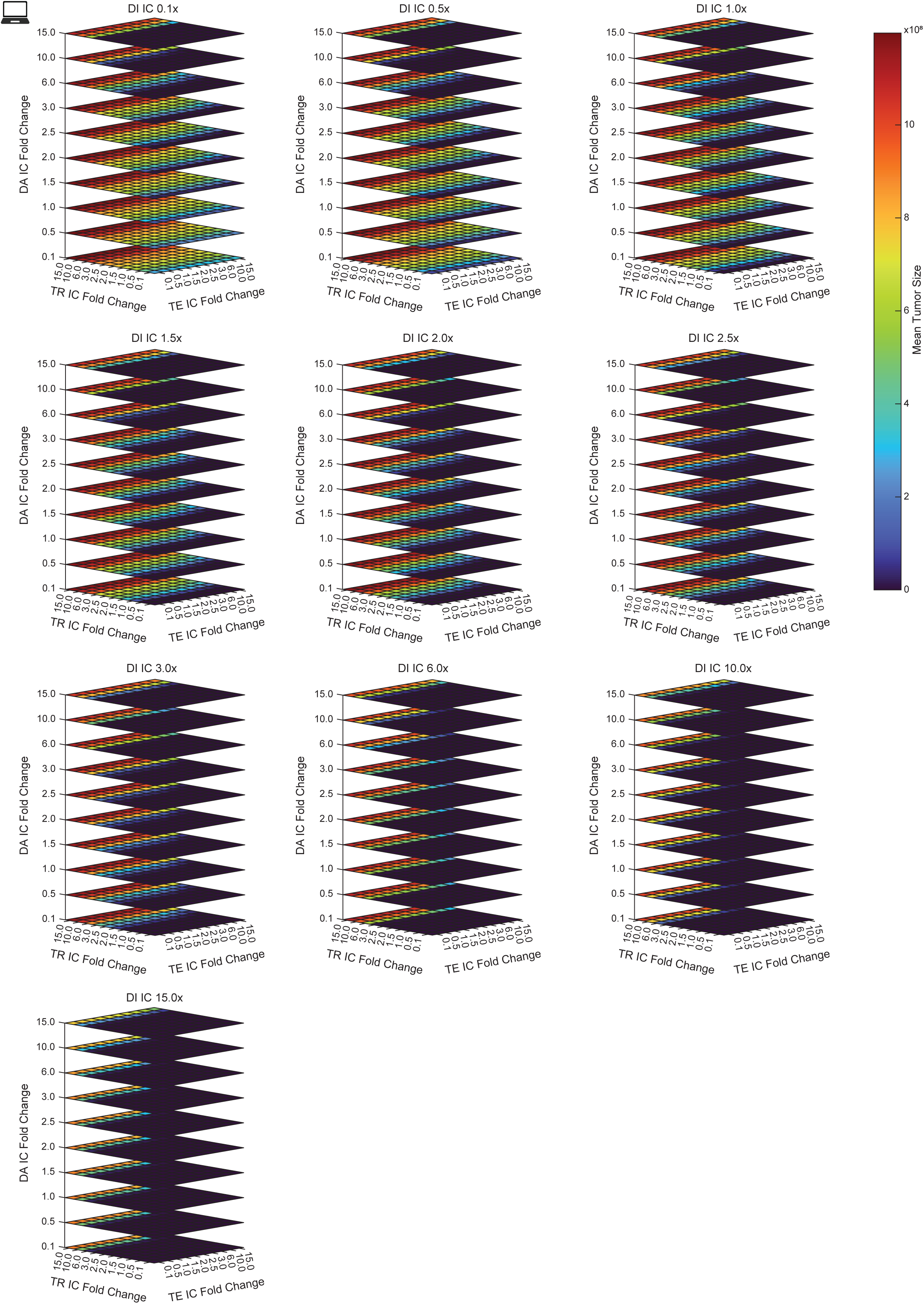
Perturbing immune cell initial conditions alters tumor outcome, related to Figure 5. Mean tumor size at final simulation time (day 35) for the indicated fold change in TE, TR, DA, and/or DI initial condition. *n* = 299 simulations/square.

**Figure S6.**
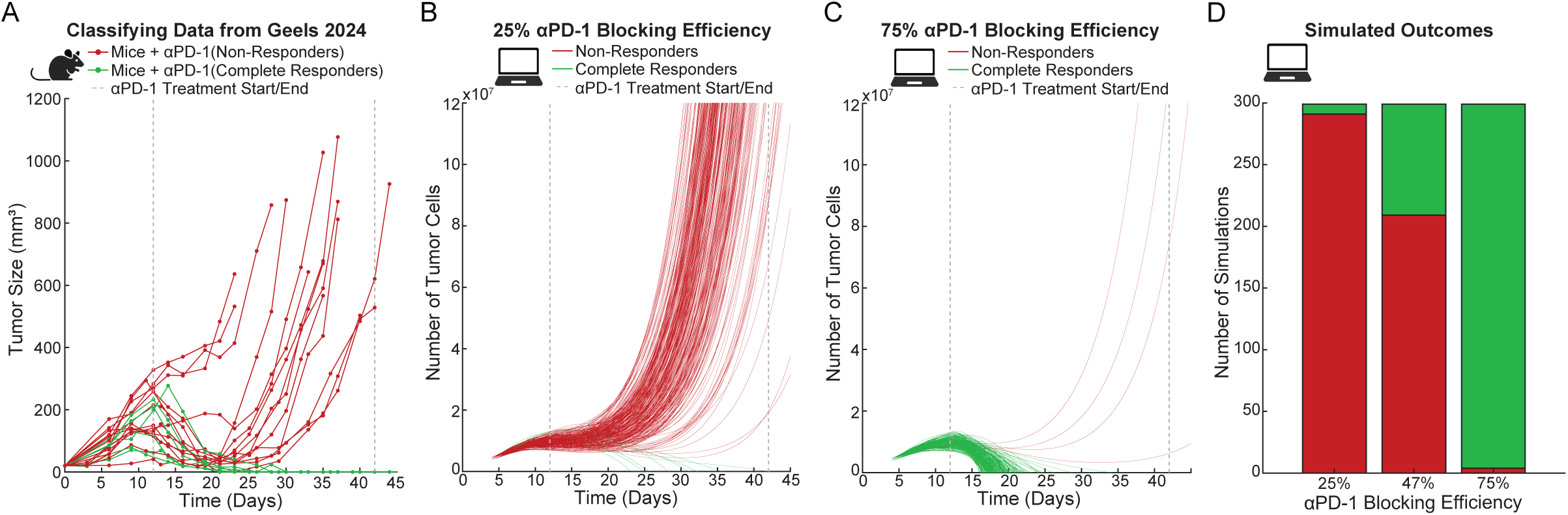
Changes in *α*PD-1 blocking efficiency alters tumor outcome, related to Figure 6. (A) Growth curves of D4M-S tumors in mice treated with *⍺*PD-1 classified as “Complete Responders” (green, tumors that are fully rejected) and “Non-Responders” (red, all other tumors). Dashed lines indicate treatment start and stop time. *n* = 20 mice from 2 separate experiments published in Geels et al., Cancer Cell 2024. (B) Simulated response to *⍺*PD-1 treatment with blocking efficiency set to 25% classified as “Non-Responders” (red) and “Complete Responders” (green). Dashed lines indicate treatment start and stop time. *n* = 299 simulations. (C) Simulated response to *⍺*PD-1 treatment with blocking efficiency set to 75% classified as “Non-Responders” (red) and “Complete Responders” (green). Dashed lines indicate treatment start and stop time. *n* = 299 simulations. (D) Simulated outcome to *⍺*PD-1 treatment by response type for three different *⍺*PD-1 blocking efficiencies. *n* = 299 simulations/group.

**Figure S7.**
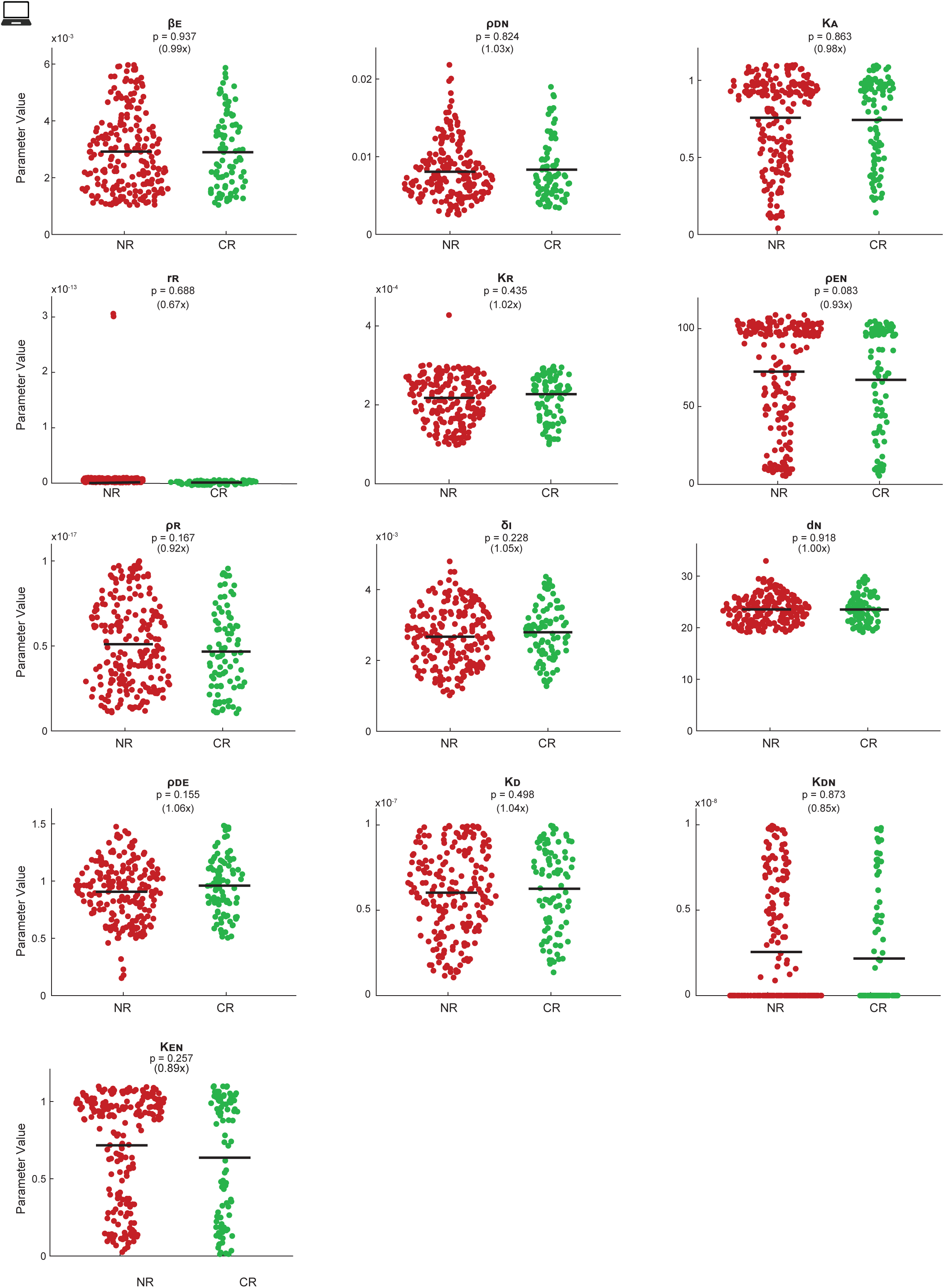
Analysis of non-significantly different parameters in responder and non-responder virtual mice, related to Figure 7. Difference between parameter values in simulations classified as “Non-Responders” (red) and “Complete Responders” (green) with simulated *⍺*PD-1 treatment. Black bars depict mean; *n* = 299 simulations; *p* values determined by Mann-Whitney U test.

## METHODS

### KEY RESOURCES TABLE

| REAGENT or RESOURCE | SOURCE | IDENTIFIER |
| --- | --- | --- |
| **Antibodies** |  |  |
| CD8α (clone: 2.43) | BioXCell | Cat# BE0061 |
| PD-1 (clone: 29F.1A12) | lchorbio | Cat# ICH1091 |
| Rat IgG | Sigma | Cat# I8015-100MG |
| B220 PE-Cy5 (clone: RA3-6B2) | BioLegend | Cat # 103210 |
| CD11c Ax700 (clone: HL3) | BD Biosciences | Cat # 560583 |
| CD135 Biotin (clone: A2F10) | BioLegend | Cat # 135308 |
| CD25 APC (clone: PC61) | BioLegend | Cat # 102011 |
| CD28 (clone: 37.51) | BioLegend | Cat# 102116 |
| CD366 (Tim3) PE-Daz (clone: RMT3-23) | BioLegend | Cat # 119748 |
| CD3ε (clone: 145-2C11) | BioLegend | Cat# 100340 |
| CD4 BV605 (clone: RM4-5) | BioLegend | Cat # 100548 |
| CD4 BV650 (clone: RM4-5) | BioLegend | Cat # 100555 |
| CD45 BV510 (clone: 30-F11) | BioLegend | Cat # 103138 |
| CD45.1 PE-Cy (clone: A20) | BioLegend | Cat # 110729 |
| CD45.2 PerCP-Cy5.5 (clone: 104) | BioLegend | Cat # 109827 |
| CD8 BV785 (clone: 53-6.7) | BioLegend | Cat # 100750 |
| FcBlock (anti-mouse CD16/32) (clone: S17011E) | BioLegend | Cat# 156604 |
| Foxp3 BV421 (clone: FJK-16s) | Invitrogen | Cat # 404577382 |
| Gr-1 PE-Cy5 (clone: RB6-8C5) | BioLegend | Cat # 108410 |
| H-2K(b) SIINFEKL Tetramer Ax647 | NIH Tetramer Core | RRID:<br>AB_3068342 |
| H-2K<sup>b</sup> PerCP-Cy5.5 (clone: AF6-88.5) | BioLegend | Cat # 116516 |
| IA/IE BV605 (clone: M5/114.15.2) | BioLegend | Cat # 107639 |
| IFNγ PE (clone: XMG1.2) | BioLegend | Cat # 505808 |
| NK1.1 PE-Cy5 (clone: PK136) | BioLegend | Cat # 108716 |
| Streptavidin PE | BioLegend | Cat # 405203 |
| TCF1 Ax488 (clone: C63D9) | Cell Signaling<br>Technology | Cat # 6444S |
| TCRβ PE-Cy5 (clone: H57-597) | BioLegend | Cat # 109210 |
| TNF APC (clone: MP6-XT22) | BioLegend | Cat # 506308 |
| **Chemicals, peptides, and recombinant proteins** |  |  |
| Brefeldin A Solution (1,000X) | BioLegend | Cat# 420601 |
| Collagenase IV | Worthington | Cat# LS004189 |
| Diphtheria Toxin | Calbiochem | Cat# 322326 |
| DNase I | Roche | Cat#<br>04536282001 |
| Tamoxifen | Sigma | Cat# T5648 |
| **Critical commercial assays** |  |  |
| Foxp3 Transcription Factor Staining Buffer Set | Invitrogen | Cat# 00552300 |
| **List of reanalyzed datasets** |  |  |
| D4M-S growth curve: experiment 3 | Geels et al., *Cancer Cell*<br>2024 | N/A |
| D4M-S growth curve: experiment 4 | Geels et al., *Cancer Cell*<br>2024 | N/A |
| D4M-S growth curve: experiment 5 | Geels et al., *Cancer Cell*<br>2024 | N/A |
| D4M-S growth curve: experiment 6 | Geels et al., *Cancer Cell*<br>2024 | N/A |
| **Experimental models: Cell lines** |  |  |
| D4M.3A H2B SIINFEKL Cer (D4M-S) | Thorsten Mempel<br>(Massachusetts General<br>Hospital and Harvard<br>Medical School, Boston,<br>MA), Di Pilato et al. 2019 | N/A |
| **Experimental models: Organisms/strains** |  |  |
| mouse: B6.129(Cg)-Foxp3tm3(DTR/GFP)Ayr/J<br>(Foxp3<sup>DTR</sup>) | Jackson Laboratories | JAX:016958 |
| mouse: C57BL/6J (B6 Jax) | Jackson Laboratories | JAX: 000664 |
| mouse: C57BL/6J-Ptprcem6Lutzy/J (B6 CD45.1) | Jackson Laboratories | JAX: 033076 |
| **Software and algorithms** |  |  |
| FCS Express 7 | De Novo Software | https://denovosoftware.com/ |
| FlowJo 10 | BD Biosciences | https://www.flowjo.com/ |
| Prism 10 | GraphPad | https://www.graphpad.com |
| MATLAB | MathWorks | https://www.mathworks.com/ |
| Scripts for model simulation | This paper | https://github.com/ImmunoimagingLab/Sousa-BioRXiv-2025 |
| **Other** |  |  |
| AccuCheck flow cytometry counting beads | Invitrogen | Cat# PCB100 |
| Live-or-dye 750/777 | Biotium | Cat# 32008-T |

### EXPERIMENTAL MODEL AND SUBJECT DETAILS

#### Cells

D4M.3A H2B SIINFEKL Cer (D4M-S) melanoma cells were obtained from Thorsten Mempel and cultured in DMEM supplemented with 10% fetal calf serum (GeminiBio) under 37 °C / 5% CO_2_ conditions. This cell line is derived from male mice^30^ and has not been authenticated. This cell line was routinely tested for mycoplasma contamination and found negative.

#### Mice

Foxp3^DTR^, CD45.1, and C57BL/6J mice were purchased from The Jackson Laboratory. Mice were enrolled in experiments at 6-10 weeks of age. D4M-S cells are originally from male mice, making them artificially immunogenic in female recipients. Because of this, we only implanted D4M-S melanomas in male mice. Mice were bred, housed, enrolled in experiments, and euthanized according to protocols approved by the Institutional Animal Care and Use Committee (IACUC) of the University of California Irvine.

### METHOD DETAILS

#### Model parametrization

We first worked to parameterize the model by searching the literature for available data that allowed us to infer parameter values (**Table 1**). For parameters in which we were unable to determine from literature, we used the constrained optimization function ‘*fmincon*’ in MATLAB to minimize the square of the difference between the data and the model fit.

#### Data unit conversions

Since our tumor data was measured in volume (mm^3^) or mass (mg) and our mathematical model tracked number of cells over time, we converted the data from tumor volume or mass to number of tumor cells according to reference ^70^.

#### Latin hypercube sampling

To generate additional parameter sets (‘virtual mice’) to use for analysis, we specified a range for each parameter of interest based on the parameter values obtained via model parameterization. We then performed Latin hyper-cube sampling using the ‘*lhsdesign’* function in MATLAB and selected parameter sets that fell within the standard deviation of the last time point for each cell type.

#### Antibody treatment of tumor-bearing mice

Mice received a subcutaneous injection of 2×10^6^ or, in the indicated experiments, of 2×10^5^ D4M-S cells. For studies ending in flow cytometry analysis, we injected two tumors per mouse 1cm off the midline on the abdomen. On day twelve, we began treatment with 200 µg αPD-1 (29F.1A12) i.p. and repeated injections every other day. We injected 200µg rat IgG i.p. as an isotype control. Tumor growth after CD8 depletion was conducted as per Ref.^10^. Briefly, 8-day-old D4M-S tumors were treated with FTY720 (1mg/kg i.p. every other day) and αCD8 or rat IgG antibodies (300 μg i.p. every third day). Tumor weight was determined six days after treatment started.

#### Measurement of immunotherapy-treated tumors

For tumor growth studies, we implanted only one tumor and measured three times a week with an electronic caliper. Tumor volume was estimated using the formula 0.5 x a x b^2^, where a is the maximum and b is the perpendicular tumor diameter. Immunotherapy administration ended on day 40 after tumor injection. When tumors reached a maximum diameter of >15mm or both diameters >10mm, mice were sacrificed.

#### Bone marrow chimeras

We created bone marrow chimeras by irradiating mice at 750 rads (X-rays source). Irradiation doses were established to ensure engraftment of 8-10 × 10^6^ donor bone marrow cells with minimal lethality. To investigate the impact of early Treg influx on tumor growth, we reconstituted CD45.1 hosts with a 1:1 mixture of Foxp3^DTR^ and either S1pr1^f/f^ or S1pr1^f/f^UBC^creERT2^ bone marrow, gently provided by Dr. Susan Schwab (New York University). Bone marrow chimeras were enrolled in experiments two months after transplantation to ensure hematopoietic reconstitution. Chimeras were given drinking water with 40mg/ml Sulfamethoxozole and 8mg/ml Trimethoprim for 1 month after irradiation to prevent opportunistic infections. We confirmed engraftment by flow cytometry analysis of blood. Tamoxifen treatment consisted of oral gavage (15 mg in 75 µl EtOH + 425 µl corn oil) on day -5 followed by four i.p. daily injections of 2 mg (in 10 µl EtOH + 40 µl corn oil). Chimeras were injected i.p. with a 25ug/kg attack dose of diphtheria toxin (Calbiochem) on day 0 followed by a 5ug/kg dose on days 1-3 (Marangoni J. Immunol 2018). We verified induction of cre by tamoxifen and depletion of Foxp3^DTR^ Tregs by diphtheria toxin through flow cytometry analysis of blood.

#### Flow cytometry

Tumors were finely minced and digested for 30 min at 37 °C using DMEM 10% FCS with 1.5 mg/ml Collagenase IV (Worthington) and 50 U/ml DNAse I (Roche). All samples were filtered through a 40μm strainer.

We stained 8×10^6^ cells and identified dead cells with Live-or-dye 750/777 (1:1000), diluted in PBS, for 30 min at 4 °C. We determined absolute cell numbers using AccuCheck flow cytometry counting beads (Invitrogen). To decrease nonspecific Ab binding, cells were incubated with 5 µg/ml FcBlock for 10 min at 4 °C. Extracellular antibody staining was carried out at 4 °C for 20 minutes in FACS buffer (PBS, 0.5% BSA, 2 mM EDTA). Cells were then permeabilized using the Foxp3 fixation-permeabilization buffer (Invitrogen), while intracellular staining was performed at 4 °C for 30 min in Foxp3 wash buffer. We acquired the samples on a NovoCyte Quanteon flow cytometer and analyzed the data using FCS Express and FlowJo.

##### T cell and APC panel

To enumerate and identify tumor specific T cells we used αCD45, αCD8, αCD4, αFoxp3, and a SIINFEKL tetramer. To evaluate exhaustion, we stained for αCD366 (Tim3), and αTCF1. To count APCs, we used a lineage cocktail of antibodies against TCRβ, B220, Gr-1, and NK1.1, as well as αH-2K^b^ (MHC-I), αIA/IE (MHC-II), αCD135, and αCD11c. After gating for live CD45^+^ Lin^−^ IA/IE^+^ cells, DCs were gated as CD11c^+^ CD135^+^.

##### Cytokine panel

To evaluate the function of exhausted tumor T cells, we stimulated them with plate-bound αCD3ε (10 µg/ml) and αCD28 (10 μg/ml) in the presence of brefeldin A (5 μg/ml) for eight hours at 37 °C. Cells were stained with αCD45, αCD8, αCD4, αFoxp3, αCD366 (Tim3), αTCF1, αIFNγ, and αTNF antibodies.

##### Chimera reconstitution panel

To verify reconstitution in the S1PR1 chimeras, we stained ACK-lysed blood with αCD4, αCD8, αCD25, αCD45.1, αCD45.2, and identified Foxp3^DTR^ Tregs with GFP.

#### Quantification and statistical analysis

When data followed a normal distribution, the Student’s *t* test was used. Otherwise, we employed the Mann-Whitney U test. To compare survival curves, the Mantel-Cox test was used. P values less than 0.05 were considered statistically significant.

